# Rapid inversion of singleton distractor representations underlies learned attentional suppression

**DOI:** 10.1101/2025.10.08.680699

**Authors:** Ziyao Zhang, Jarrod A. Lewis-Peacock

## Abstract

In visually complex and dynamically changing environments, humans must often filter out salient but task-irrelevant stimuli. Prior work shows that with repeated exposure to color singleton distractors, individuals can learn to divert attention away from these salient items. However, the neural mechanisms supporting such attentional suppression remain unclear. The present study examined the temporal trajectories of singleton distractor representations during visual search to address this gap. Using multivariate pattern analyses of EEG data in human subjects (N = 40, 30 females, 10 males), we identified two clusters of decodable singleton distractor representations: an early cluster from 100-200 ms and a later cluster from 200-400 ms. Temporal generalization analyses showed that the later representations were inverted versions of the early ones. Importantly, stronger late, but not early representations, predicted faster search responses, suggesting that the later signals support distractor suppression. This representational inversion facilitates suppressing singleton distractors in the spatial priority map. Comparing decoding evidence across locations revealed that singleton distractor locations were suppressed relative to non-singleton distractors. Moreover, comparing the neural coding of locations revealed that the spatial organization in the singleton distractor neural space was inverted relative to that in the target neural space. Together, these findings reveal a rapid representational inversion underlying salient distractor suppression at the onset of visual search. This inversion of singleton distractor signals was likely driven by top-down control mechanisms that transform bottom-up saliency signals, producing an inverted arrangement of target and distractor information within a shared neural space.

**Significance Statement:** A primary goal of attention is to select relevant information while ignoring irrelevant input from the external environment. Although the mechanisms of attentional enhancement have been extensively studied, the mechanisms underlying attentional suppression remain less well understood. While prior work has made important progress in identifying when attentional suppression is engaged, the neural mechanisms that implement suppression are still unclear. Here, we show that neural representations of salient singleton distractors undergo a rapid inversion approximately 200 ms after search onset. These inverted representations can subsequently be read out as suppression signals during the computation of spatial priorities. Together, our findings suggest that transforming initial pop-out distractor signals into an inverted representational format might drive rapid attentional suppression and support goal-directed visual search.

## Introduction

The ability to filter distracting information is crucial for goal-directed behavior. Salient but irrelevant stimuli, such as website pop-ups, readily capture attention, raising the question of whether such stimuli can be actively suppressed within the attention system. Recent progress on this issue has come from studies of color singleton distractors (e.g., a red item among green items) in goal-directed visual search (Bacon & Egeth, 1994; Gaspelin et al., 2015). Color singletons readily capture attention and impair search performance (Jonides & Yantis, 1988; Theeuwes, 2010). However, accumulating evidence suggests that attentional capture by such distractors can be mitigated.

Repeated exposure to search arrays containing color singleton distractors can lead to faster detection of shape-defined targets when a singleton is present than when it is absent (Chang & Egeth, 2019; Gaspelin et al., 2015, Gaspelin et al., 2017; Vatterott & Vecera, 2012). Although this behavioral benefit has been interpreted as evidence for attentional suppression, alternative mechanisms could also account for these effects, including ignoring singletons in a serial search mode (Theeuwes, 2023), down-weighting the distractor feature dimension (Liesefeld & Müller, 2019, 2021), or rapid rejection of singleton distractors (Liesefeld et al., 2021, 2025).

Eye-tracking studies show that initial saccades are less likely to land on singleton distractors than on non-singleton distractors (Gaspelin et al., 2017; Stilwell et al., 2023), suggesting suppression in overt attention. However, oculomotor measures cannot rule out rapid covert capture or fast reactive mechanisms that resolve distractor interference (e.g., Klink et al., 2023). Converging electrophysiological evidence has identified the distractor positivity component (Pd), emerging approximately 100 - 300 ms after stimulus onset in conditions associated with behavioral suppression (Cosman et al., 2018; Gaspelin et al., 2023; Sawaki & Luck, 2010; Sawaki et al., 2012). This suggests that the suppression might be associated with early inhibitory processes. However, the functional role of Pd remains debated, with alternative accounts proposing that it reflects upweighting of non-distractor representations (Kerzel & Burra, 2020; Kerzel & Huynh Cong, 2023) or sensory imbalances between visual hemifields (Oxner et al., 2025).

These findings have motivated the signal suppression hypothesis, which posits that salient stimuli automatically generate bottom-up priority signals that can be suppressed by top-down mechanisms before spatial attention is deployed (Gaspelin & Luck, 2018b; Sawaki & Luck, 2010; Sawaki et al., 2012). In this framework, suppression operates by down-weighting the distractor feature dimension prior to the initial attentional shift (Gaspelin et al., 2018b, 2025). In contrast, the rapid disengagement hypothesis proposes that suppression occurs reactively, after attention has been captured by the distractor (Theeuwes, 2010; Theeuwes et al., 2000). Recent computational modeling suggests that these accounts may reflect differences in the relative timing of target- and distractor-related priority signals rather than distinct mechanisms (Zhang et al., 2025).

Despite advances in behavioral and computational accounts, the neural mechanisms underlying attentional suppression remain unclear. One possibility is neural suppression, whereby distractor-related activity or representations are reduced during visual search (Turatto et al., 2018; Vatterott & Vecera, 2012; Won et al., 2019). Consistent with this account, fMRI and non-human primate studies have reported reduced neural responses to, and weakened representations of, repeated singleton distractors in early visual cortex and in priority-map regions, including intraparietal cortex, frontal eye fields (FEF), and lateral intraparietal cortex (LIP) (Adam & Serences, 2021; Cosman et al., 2018; Ipata et al., 2006; Won et al., 2020).

In contrast, recent evidence suggests that attentional suppression may rely on mechanisms other than uniform neural suppression. Neuronal populations in V4, FEF, and LIP exhibit heterogeneous response profiles to singleton distractors, rather than consistent reductions in activity (Klink et al., 2023; Sapountzis et al., 2025). Moreover, decoding analyses indicate that distractor representations remain stable with training, rather than being eliminated or suppressed, raising the possibility that these representations are coded in a format that facilitates rather than interferes with attentional allocation to targets (Sapountzis et al., 2025).

Here, we tested and provided evidence for representational transformation of singleton distractors during visual search using temporal generalization analyses of EEG data. Whereas neural suppression accounts predict that distractor representations weaken over time, transformation accounts predict changes in representational format without necessarily implying reduced representational strength. Consistent with the latter, we observed reliable representations of singleton distractor locations across time. Notably, the representational format transformed approximately 200 ms after search onset; specifically, later representations reflected an inverted version of the early representations.

## Materials and Methods

We reanalyzed data shared from Stilwell et al., 2022. Two groups of 20 participants were recruited for experiment 1 and experiment 2, respectively (N = 40, 30 females, 10 males). In each trial, participants searched for a predefined target item (circle or diamond, counterbalanced between participants) in a search array. In the search array, the inner ring contained a target shape and distractors that were in different shapes. On 75% of trials, a color singleton distractor that had a distinct color from other items was presented in the inner ring. The color singleton distractor was never the target, and participants were instructed to use the shape feature to locate the target item. The specific color of the color singleton distractor (red or green) was fixed for each participant but counterbalanced across participants. The outer ring contained non-target shapes only to boost the relative salience of the singleton distractor. Overall, participants completed 1296 singleton-present trials and 432 singleton-absent trials. Search arrays in experiment 1 had an inner ring with four items (effective set size 4), while in experiment 2, the inner ring had eight items (effective set size 8). EEG signals were collected and preprocessed following the procedures outlined in Stilwell et al., 2022.

### Decoding of target and singleton distractor locations

Location decoding was conducted separately for singleton and singleton-absent trials. For singleton-absent trials, we used support vector machine (SVM) to classify the target item’s location based on the spatial distribution of the EEG signal across 17 posterior electrodes (Pz, P3, P5, P7, P9, PO7, PO3, O1, POz, Oz, P4, P6, P8, P10, PO4, PO8, O2). We implemented this model using the *svc* function from the *sklearn* package in Python. EEG data were downsampled to 100 Hz, and decoding was performed for each time point (10 ms) ranging from -200 ms to 800 ms relative to the onset of the search array. For each time point, the dataset was divided into 3 folds. An equal number of trials from each location class were randomly selected for each fold without replacement to ensure unbiased training. Two folds were used as the training set, and the remaining fold served as the testing set. This procedure was repeated for the 3 folds, with each one serving as the testing set in an iteration. The entire process was repeated 100 times (Wolff et al., 2020), and decoding accuracies were averaged across 100 iterations. Decoding of target and singleton distractor locations in singleton-present trials followed the same procedure as decoding target locations in singleton-absent trials, except for adjusting the training set to match the number of trials in singleton-absent trials. This adjustment aimed to ensure equal training data for decoders in both singleton and singleton-absent trials. This should prevent biases in decoding performance comparisons between singleton and singleton-absent trials resulting from uneven training sets.

#### Linear models linking decoding evidence to reaction time

To link target location representations to reaction time, decoding evidence for the correct target location in each trial was obtained from the *decision_function* in *sklearn*, which reflects the model’s confidence (distance to the decision hyperplane). Higher values indicate stronger neural representations of the target location. Decoding evidence was averaged within the early (100 - 200 ms) and later (200 - 400 ms) time windows. Both the averaged decoding values and reaction times were z-scored within participants before fitting the linear models. Linear mixed effects models were then fitted using the *lmer* function in *R*, with early and late decoding evidence as predictors and reaction time as the outcome variable. To link singleton distractor representations to reaction time, we applied the same analysis pipeline, except that decoding evidence was extracted for singleton distractor locations rather than target locations.

#### Latency estimation

To estimate the onset latency of above-chance decoding, we applied a jackknife-based bootstrapping procedure (Miller et al., 1998; Ulrich & Miller, 2001). We generated N bootstrapped subsamples (N = 20 in both experiments), each created by leaving out one participant. For each subsample, we performed a one-sample t-test comparing decoding accuracy to chance. The latency for that subsample was defined as the onset of the earliest cluster of significant time points. This procedure yielded N latency estimates. These estimates were then used for statistical comparisons.

### Temporal generalization analysis

Temporal generalization analyses were conducted for target locations in singleton-absent trials and singleton locations in singleton-present trials. The same decoding procedures were used, with the only difference being that trained decoders for each time point were used to predict location labels for data at all time points.

### Cross-condition generalization

Cross-condition generalization was performed between target locations in singleton-absent trials and singleton distractor locations in singleton-present trials. SVM decoders were trained exclusively on data from singleton-absent trials with labels indicating target locations. These trained decoders were then used to predict singleton distractor locations in singleton-present trials. Decoding accuracies were computed based on the probability of the decoders’ predictions matching the actual singleton distractor location labels.

### Activation scores

To compute activation scores for each location in singleton-present trials, decoders were first trained on target locations in singleton-absent trials. These trained decoders were subsequently used to predict location labels in singleton-present trials. Instead of generating prediction labels, prediction probabilities were produced to indicate the likelihood of a given location being the true label based on the neural patterns. Prediction probabilities for target locations, singleton locations, and non-singleton distractor locations were averaged across trials for each subject. For plotting the spatial priority map, the averaged prediction probabilities of non-singleton distractor locations (two in experiment 1, six in experiment 2) were used as the baseline activation. The activation for target and singleton distractor locations was computed as the difference between the prediction probabilities of these locations and the baseline.

### Correlations between target and singleton representations

Raw EEG data of singleton-absent trials (200 - 400 ms) with the same target location were averaged to create the mean target representations for each potential target location (four in experiment 1 and eight in experiment 2). Likewise, singleton-present trials with the same singleton distractor location were averaged (200 - 400 ms) to generate singleton distractor representations for all potential singleton distractor locations (four in experiment 1 and eight in experiment 2). Pearson correlations in multi-channel EEG activities between target and singleton representations were computed for each potential location and then averaged. Positive correlations indicate similarity between the location representations in the target space and location representations in the singleton distractor space, while negative correlations suggest inverted representations.

Principal Component Analysis (PCA) was utilized to visualize the relation between the target representational space and the singleton distractor representational space. For this purpose, we identified the first two principal components (PCs) of the target representational space for one example participant (99% variance explained). Both target and singleton distractor representations were projected into the space defined by these two PCs. It should be noted that PCA was used to visualize the relationship between the target and singleton distractor representation only. The formal correlation analysis was performed based on the raw signal space.

### Statistics

We conducted one-sample (one-sided) t-tests to compare decoding accuracies against the theoretical chance level, as below-chance decoding is not meaningful for this analysis. All other comparisons were performed using two-sided t-tests. One-sample t-tests were also used to compare temporal generalizability scores and cross-condition generalizability scores against the theoretical chance level. In both cases, above-chance scores indicated positive generalizability, whereas below-chance scores indicated negative generalizability. Paired t-tests were used to compare prediction probabilities for target locations, singleton distractor locations, and non-singleton distractor locations. Finally, one-sample t-tests were conducted to compare neural correlations between target and singleton distractor representations against zero, with positive values indicating positive correlations and negative values indicating negative correlations. All analyses were performed in Python using standard statistical packages. Bayes factors for the alternative hypothesis (*BF10*) were reported, with values between 1 and 3 indicating anecdotal evidence, values between 3 and 10 indicating moderate evidence, and values greater than 10 indicating strong evidence. To account for multiple comparisons, Sidak correction was applied, and adjusted p-values were reported.

## Results

Participants searched for a predefined target item (circle or diamond, counterbalanced between participants) in a search array. In the search array, the inner ring contained a target shape and distractors that were in different shapes. On 75% of trials, a color singleton distractor that had a distinct color from other items was presented in the inner ring. The color singleton distractor was never the target, and participants were instructed to use the shape feature to locate the target item. The outer ring contained non-target shapes only to boost the relative salience of the singleton distractor.

### Singleton distractor representations showed distinct temporal patterns compared to target representations

We applied MVPA to track target and singleton distractor representations across time. Both singleton distractor and target locations showed successful decoding (**Fig 1**). Target location representations emerged around 200 ms and persisted for most of the search period. Interestingly, the decoding of singleton distractor locations revealed an earlier initial peak, spanning from 100 ms to 200 ms. This initial peak was followed by a subsequent, more robust peak occurring between 200 ms and 400 ms for both set size 4, and set size 8. The decoding accuracies of singleton distractor locations quickly dropped off following the second peak whereas that for target locations persisted. The estimated latency for singleton distractor representations was faster than that for target representations in both singleton-present trials [experiment 1: *t*(19) = -5.17, *p* < 0.001, *d* = 1.58, *BF10* = 477.01; experiment 2: *t*(19) = -259.0, *p* < 0.001, *d* = 81.90, *BF10* = *1.2 × 10^31^*] and singleton-absent trials [experiment 1: *t*(19)= -9.92, *p* < 0.001, *d* = 3.15, *BF10* = *2.0 × 10^6^*; experiment 2: *t*(19) = -46.15, *p* < 0.001, *d* =14.59, *BF10* = *4.3 × 10^17^*]. Interestingly, the latency for target representations in singleton-present trials was faster than in singleton-absent trials [experiment 1: *t*(19) = -11.86, *p* < 0.001, *d*= 3.87, *BF10* = *3.1 × 10^7^*; experiment 2: *t*(19) = -12.11, *p* < 0.001, *d* = 3.79, *BF10* = *4.4 × 10^7^*].

**Fig. 1.**
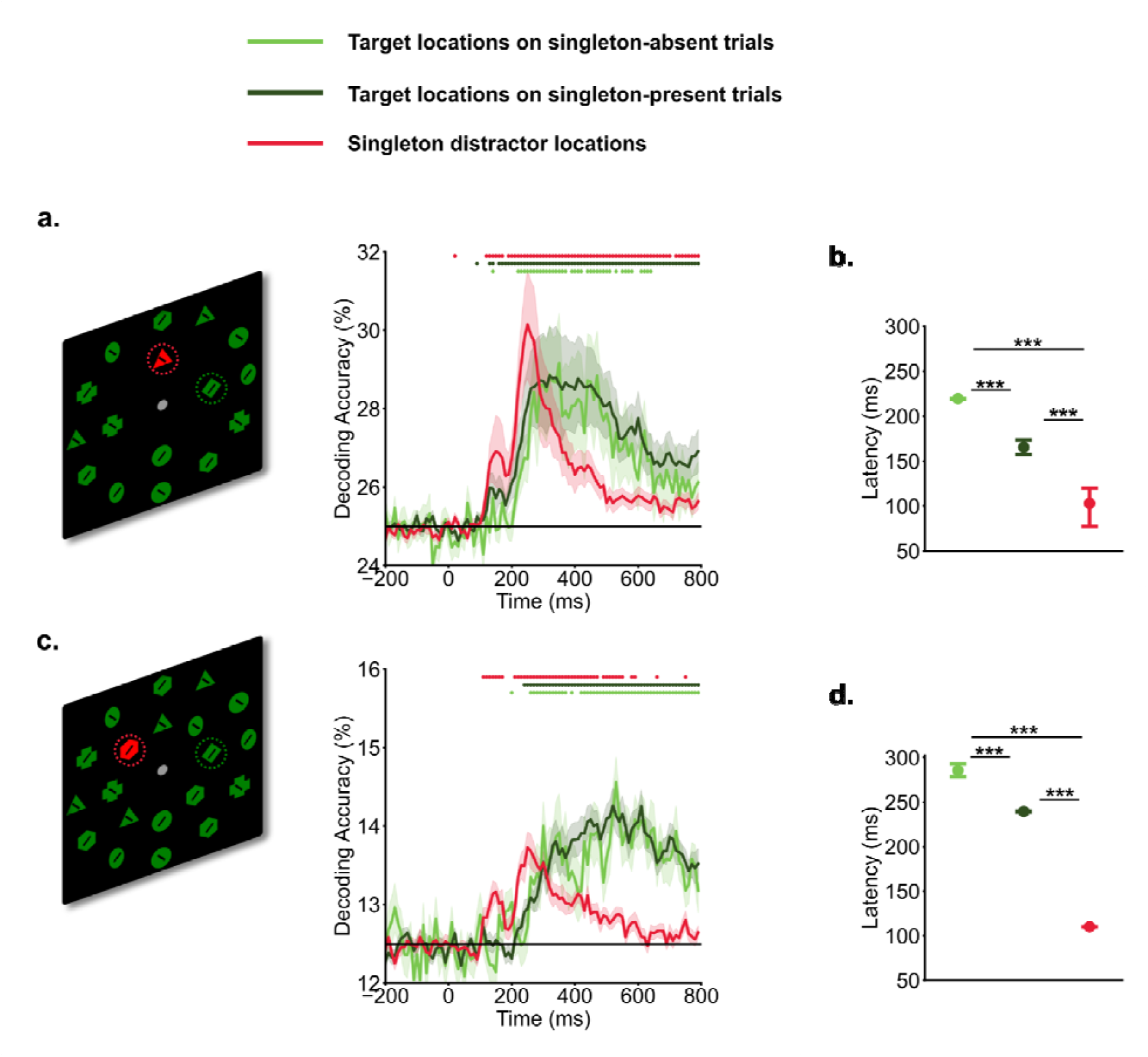
Temporal profiles of target and singleton distractor representations. Participants searched for a specific target shape (e.g., a green diamond) within the inner ring of a search array, and made a speeded button press to indicate the direction of the line within the target shape (left vs. right). Time 0 ms represents the onset of search arrays. In experiment 1 (a. set size 4) and experiment 2 (b. set size 8). a&c). decoding of singleton distractors showed an earlier initial peak of decoding evidence from 100 ms to 200 ms, followed by a later peak from 200 ms to 400 ms. Target decoding showed a gradual increase in evidence The horizontal black line indicates chance-level (25% for se size 4, and 12.5% for set size 8). Shaded areas indicate standard errors. Colored dots indicate above chance-level decoding performance (*p* < .05, one-sided t test). b&d). Latency of decoding accuracies estimated via the jackknife-based procedure. *** indicates p < .001.

The identification of two early peaks in singleton distractor decodings suggests potential shifts in representation during the search process. We conducted temporal generalization analyses to further examine these potential changes. Our methodology involved training decoders at one time point and then testing the decoder across all time points, repeating this procedure for each time point. As shown in **Fig 2**, target representations were stable over time. Trained decoders were able to generalize to neighboring time points. In contrast, the temporal generalization analysis of singleton distractor representations showed two discernible clusters. The initial cluster extended approximately from 100 ms to 200 ms, while the subsequent cluster emerged at around 200 ms. Crucially, these two clusters showed an inverted relationship. Decoders trained on activity from 100 ms to 200 ms had below-chance decoding performance when tested on data from 200 ms to 400 ms and vice versa. This negative generalizability performance suggests an inversion of representations for singleton distractors between the early (100 - 200 ms) and the later (200 - 400 ms) time windows. It also provides preliminary evidence for the signal transformation hypothesis that neural codings of singleton distractors were inverted during the search. However, it is unclear yet how this inversion directly supported attentional suppression.

**Fig. 2.**
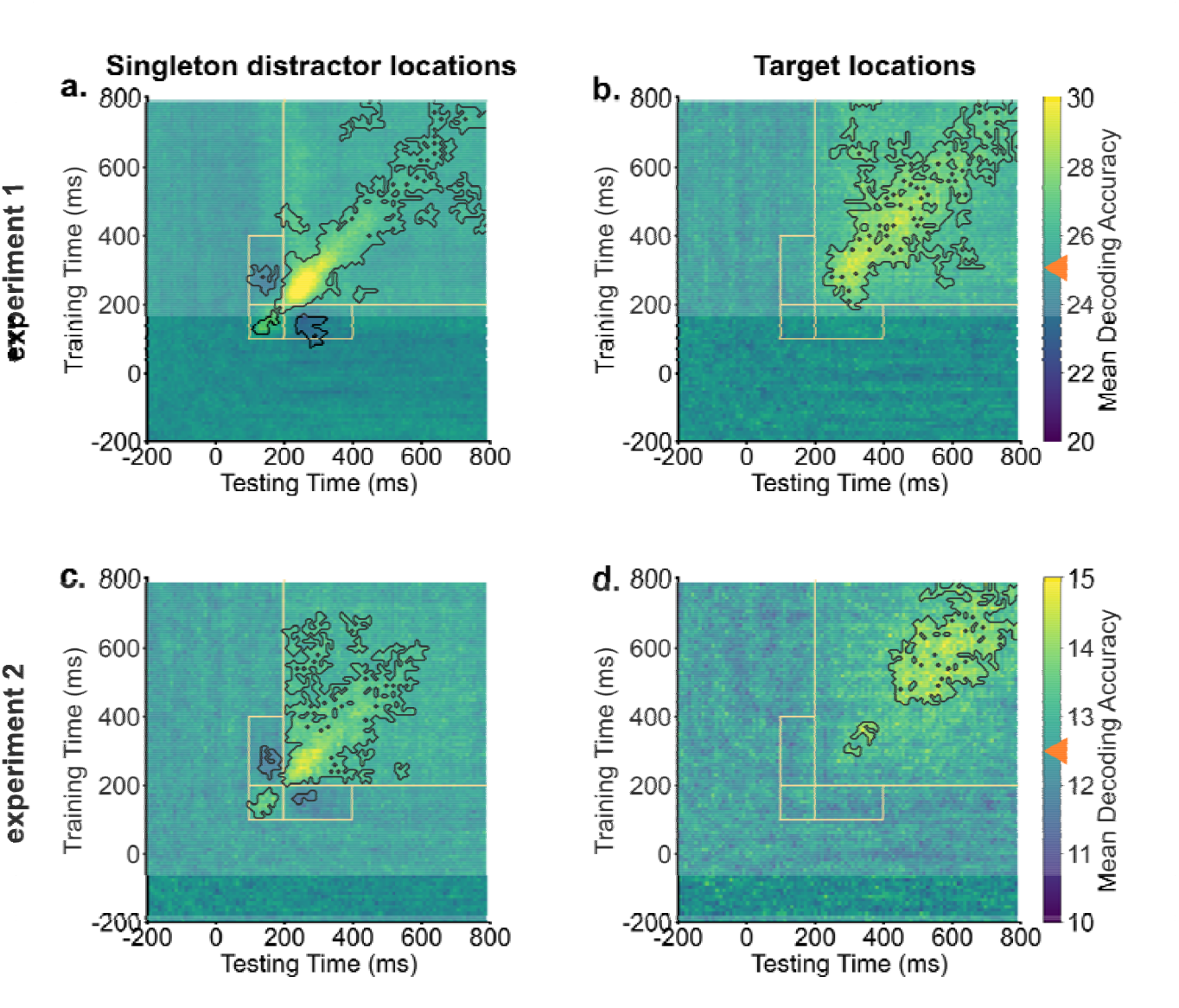
Singleton distractor representations showed inverted transformations, whereas target representations remained stable. Across experiments, generalization analyses of singleton distractor representations (a & c) revealed two clusters. Training decoders on activities from 100 ms to 200 ms resulted in below chance-level decoding evidence when tested on activities from 200 ms to 400 ms, and vice versa. Target representations demonstrated stability and could be generalized to neighboring time points (b & d). Orange boxes indicate key generalization time windows. Black contours indicate significant areas (*p* < .05). Bright colors indicate above chance-level decoding performance (denoted by orange triangles), and dark colors indicate below chance-level decoding performance.

Singleton distractor representations facilitated visual search. We linked trial-by-trial decoding evidence for target and singleton distractor locations to reaction time to examine their functional roles. The relationship between early (100 - 200 ms) and late (200 - 400 ms) singleton distractor representations and reaction time is particularly informative for distinguishing between theoretical accounts of distractor processing. Under the reactive suppression (rapid disengagement) framework, attentional suppression occurs after attention has been captured by the singleton. Accordingly, early distractor representations would be expected to reflect attentional capture, whereas later representations would reflect reactive suppression. Based on this account, early distractor representations should be associated with slower reaction times, whereas later representations should be associated with faster reaction times. In contrast, under the proactive suppression framework (signal suppression), both early and late distractor representations likely reflect different stages of active suppression. Under this framework, both representations, particularly the later ones, should be associated with faster reaction times. Our linear modeling results (Fig 3) showed that only late target representations negatively predicted reaction times in both singleton-absent (experiment 1: β = -0.04, CI = [-0.06, -0.02], *p* = .001, *BF10* = 11.22; experiment 2: β = -0.06, CI = [-0.08, -0.03], *p* < .001, *BF10* = 76.21; combined: β<ent>= -0.05, CI = [-0.06, -0.03], *p* < .001, *BF10* = 21555) and singleton-present trials (experiment 1:β = -0.07, CI = [-0.08, -0.05], *p* < .001, *BF10* = *2.3 × 10^20^*; experiment 2: β = -0.08, CI = [-0.10, -0.06], *p* < .001, *BF10* = *2.1 × 10^29^*; combined: β = -0.07, CI = [-0.08, -0.06], *p* < .001, *BF10* = *6.7 × 10^50^*), whereas early target representations did not ( *ps* > .05). Critically, early singleton distractor representations did not predict reaction times (experiment 1: β = 0.00, CI = [-0.01, 0.01], *p* = .942, *BF10* = 0.009; experiment 2: β = 0.01, CI = [-0.00, 0.03], *p* = .083, *BF10* =0.021; combined: β = 0.01, CI = [-0.00, 0.02], *p* = .199; *BF10* = 0.011), whereas late distractor representations negatively predicted reaction times (experiment 1: β = -0.02, CI = [-0.03, 0.00], *p* = .008, *BF10* = 65.79; experiment 2: β = -0.02, CI = [-0.03, -0.00], *p* = .035, *BF10* = 3.52; combined: β = -0.02, CI = [-0.03, -0.01], *p* = .001, *BF10* = 11.45). Together, these findings indicate that both target and singleton-distractor representations in the late window are associated with faster reaction times and thus enhanced behavioral performance. Distractor representations in the early time window did not predict behavioral performance, which runs counter to a rapid disengagement interpretation of our results. The signal suppression hypothesis could potentially account for this pattern, but only if the early component of distractor processing reflects an initial registration of salience that is subsequently used for diverting attention away. However, this interpretation would be less consistent with a strict proactive suppression framework, which posits that suppression is implemented prior to the onset of the search array.

**Fig. 3.**
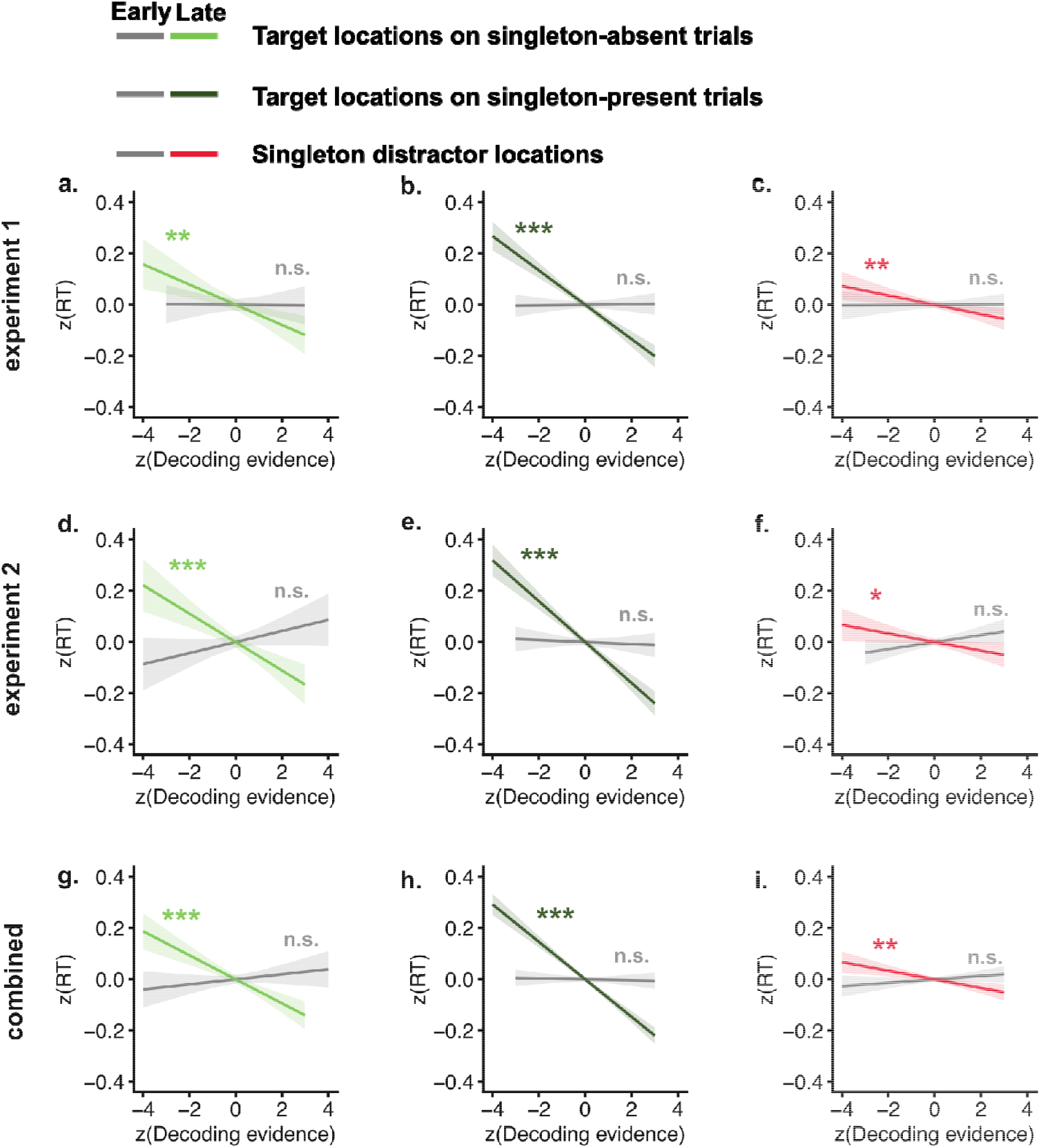
Target and singleton distractor representations predict faster reaction times. a–c) Target representations in singleton-absent, singleton-present, and singleton distractor representations predicted reaction time in experiment 1. Gray indicates the early window (100 - 200 ms); light green, dark green, and red indicate the late window (200 - 400 ms). d–f) Linear model results for experiment 2. g–i) Linear model results combining experiments 1 and 2. Shaded areas represent confidence intervals. n.s indicates non-significant. * indicates p<.05, ** indicates p < .01, *** indicates p < .001.

### Singleton distractor locations were suppressed in the spatial priority map

To investigate how the inversion of singleton representations supported attentional suppression and facilitated visual search, we performed cross-condition generalization analyses. In this procedure, decoders were trained on target representations in singleton-absent trials, where top-down enhancements served as the primary guiding sources to boost target locations in the priority map. Subsequently, these trained decoders were tested on singleton-present trials to reveal the priority maps in instances where both target enhancement and singleton suppression might influence spatial priorities. The cross-condition generalization analyses revealed that singleton distractor locations were suppressed in the spatial priority map, resulting in below chance-level decoding performance (**Fig 4.a & b**). This suppression of singleton distractor locations started around 200 ms and persisted. Intriguingly, no generalization was observed between target representations and singleton distractor locations during the early time window from 100 ms to 200 ms, corresponding to the initial cluster of singleton distractor representation observed in temporal generalization analyses.

**Fig. 4.**
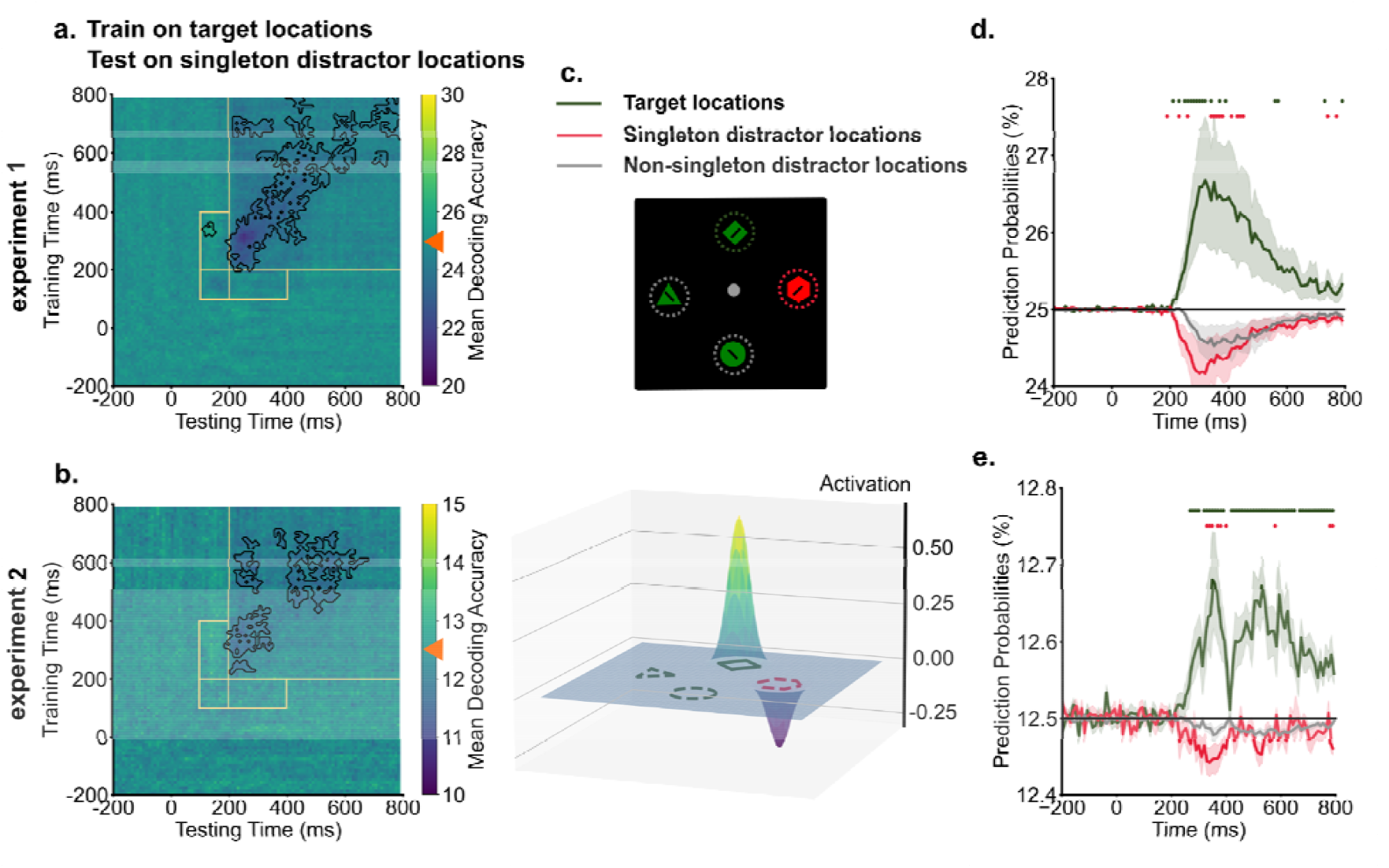
Singleton distractor locations were suppressed in the spatial priority map. a & b) Across experiments, training decoders on target locations from singleton-absent trials demonstrated below chance-level decoding performance (denoted by orange triangles) for singleton distractor locations, indicating the suppression of singleton distractor locations. Dark colors indicate below chance level decoding performance for singleton locations. Black contours indicate significant areas (*p* < .05). c) An example (250 ms in experiment 1) illustrates how we computed activation scores from decoding prediction probabilities. Prediction probabilities of target locations (green) and singleton distractor locations (red) were baselined to the averaged prediction probabilities of non-singleton distractor locations (gray). d & e) The time course of prediction probabilities for target locations and singleton distractor locations were plotted against the baseline (other locations). From 200 ms into the search, target locations showed increased prediction probabilities, whereas singleton distractor locations showed decreased prediction probabilities compared to the baseline. Shaded areas indicate standard errors. Colored dots indicate significant differences in prediction probabilities between target, singleton distractor locations to non-singleton distractor locations (*p* < .05).

Below-chance decoding performance could reflect suppression of singleton distractor locations. Alternatively, it might be driven by a strong enhancement of target locations, which would reduce decoding performance for all other locations. To rule out the alternative possibility, we compared the spatial priority of singleton distractors against a baseline of non-singleton distractors in the search array. We derived prediction probabilities from trained decoders for each location in the search array and compared them across target locations, singleton distractor locations, and non-singleton distractor locations. Activation scores for target and singleton distractor locations were computed as the difference in prediction probabilities compared to the baseline (non-singleton distractor locations, for one example time point, see **Fig 4.c**). For between 200 ms and 400 ms into the search, we found enhancement of target locations compared to the baseline [experiment 1: *t*(19) = 2.14, *p* = 0.045, *d* = 0.81, *BF10* = 1.51; experiment 2: *t*(19) = 2.93, *p* = 0.009, *d* = 1.02, *BF10* = 5.78]. Critically, suppression of singleton distractor locations compared to the baseline was also observed across the experiment [experiment 1: *t*(19) = -2.11, *p* = 0.048, *d* = 0.32, *BF10* = 1.42; experiment 2: *t*(19) = -2.62, *p* = 0.017, *d* = 0.68, *BF10* = 3.30]. As shown in **Fig 4.d & e**, prediction probabilities of target locations displayed enhancement relative to the baseline from 200 ms onwards. In contrast, prediction probabilities of singleton distractor locations were suppressed below the baseline.

### Singleton distractor and target locations were coded in a shared neural space but in a reversed manner

Cross-condition generalization analyses suggested that singleton distractors were suppressed in the neural space representing target locations. We hypothesized that this suppression might be driven by coding distractor information in an inverted format relative to target information. Such inverted coding of target and distractor information could facilitate the integration of top-down enhancement and suppression in computing spatial priority (Awh et al., 2012; Fecteau & Munoz, 2006; Itti & Koch, 2000; Luck et al., 2021). To directly test this hypothesis, we conducted neural correlation analyses to examine the relationship between target and singleton distractor representations.

In **Fig 5.a**, representations of both targets and singletons were shown for an example subject from 200 ms to 400 ms into the search, projected in a 2D subspace for easier visualization. This subspace was constructed using the first two principal components (a total of 99% variance explained) of the neural representations of target locations. The location information demonstrated clear organization; the responses to four locations were well separated, and the distance between neural representations corresponded closely to the physical distance between locations (i.e., neighboring locations had neighboring representations). Interestingly, singleton distractor and target representations showed a reversed pattern. For instance, the top location (blue square) in the target representational space was projected to the top-right corner, whereas the same location in the singleton distractor representational space was projected to the bottom-left region (blue circle).

**Fig. 5.**
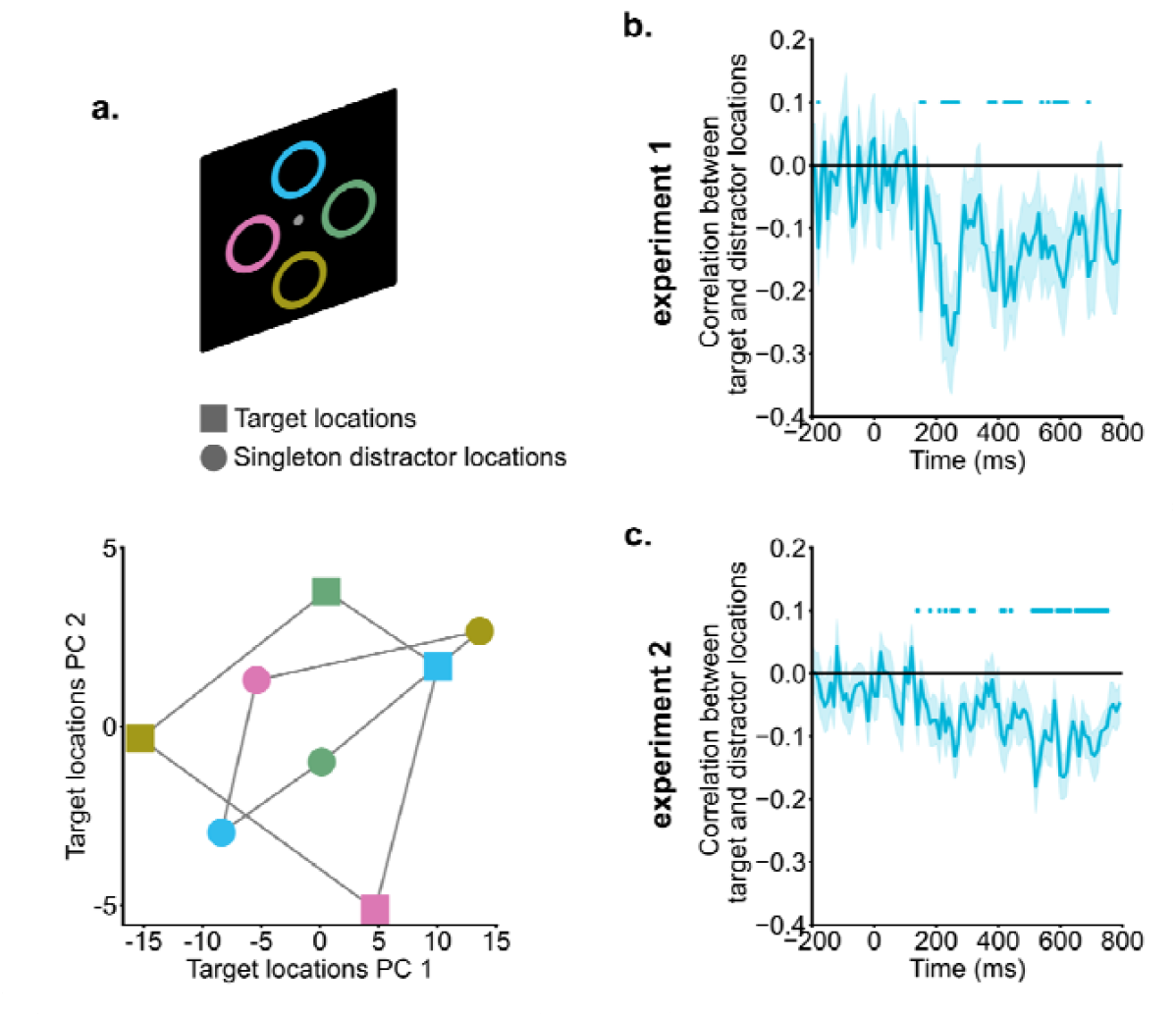
Singleton distractor suppression and target enhancement relied on inverted neural coding. a) Singleton distractor and target representations of an example subject were projected onto a 2D target representational space for illustration purposes. Different colors indicate different locations. Singleton distractor and target representations exhibited an inverted pattern. b & c) Correlations between singleton distractor and target representations. Starting from around 200 ms, singleton distractor and target representations showed negative correlations. Shaded areas indicate standard errors. Colored dots indicate significant correlations compared to zero (*p* < .05).

Formal correlation analyses were performed to link target representations and singleton distractor representations in the raw signal space (dimension = channel). For between 200 ms and 400 ms into the search, target and singleton distractor representations showed reliable negative correlations [experiment 1: *t*(19) = -3.22, *p* = 0.005, *d* = 0.72, *BF10* = 9.99; experiment 2: *t*(19) = -3.05, *p* = 0.006, *d* = 0.68, *BF10* = 7.29]. As shown in **Fig 5.b & c**, negative correlation emerged around 200 ms and persisted into the later search period, suggesting that singleton distractor representations and target representations were coded with an inverted representational geometry.

## Discussion

Recent attention research has increasingly shifted from an exclusive focus on target enhancement to a broader consideration of distractor suppression (Geng et al., 2019; Luck et al., 2021; Theeuwes, 2025; Wöstmann et al., 2022). Although converging evidence indicates that attention can be directed away from cued or learned distractors (Arita et al., 2012; Chang & Egeth, 2019; Gaspelin et al., 2015, Gaspelin et al., 2017; Sawaki & Luck, 2010; Vatterott & Vecera, 2012; Won et al., 2019; Zhang et al., 2020, 2022; Zhang & Carlisle, 2023), it remains unclear how suppression unfolds over time and how to-be-suppressed information is represented neurally. The primary goal of the present study was to test a signal transformation mechanism underlying singleton distractor suppression during visual search. Using multivariate pattern analysis (MVPA) of EEG data, we characterized the temporal evolution of singleton distractor representations.

We found that singleton distractor representations emerged reliably earlier than target representations, consistent with the idea that salient distractors generate rapid bottom-up signals prior to the emergence of top-down target enhancement (Jonides & Yantis, 1988; Theeuwes, 2010). Increasing set size from four to eight items delayed the emergence of target representations but did not affect singleton distractor representations, in line with prior work showing that crowding weakens target signals (Reddy & VanRullen, 2007) while salience-driven signals remain relatively stable. Interestingly, target representations emerged earlier in singleton-present than singleton-absent trials. Although this pattern could suggest facilitation of target processing by the presence of a singleton distractor, it may also reflect a limitation of spatial decoding in displays where target and singleton locations are mutually exclusive, as discussed below.

Decoding analyses revealed two distinct peaks in singleton distractor representations. Temporal generalization analyses showed that target representations were stable across time, forming a sustained cluster from approximately 200 ms onward. In contrast, singleton distractor representations formed two temporally dissociable clusters: an early cluster from 100 - 200 ms and a later cluster around 200 - 400 ms. Critically, these two singleton clusters exhibited negative cross-temporal generalization, indicating a representational inversion around 200 ms. This pattern provides direct evidence that singleton distractor representations undergo a transformation rather than a monotonic attenuation.

To assess how this representational inversion influenced spatial priority computations, we trained decoders on target locations from singleton-absent trials, where spatial priority is primarily driven by top-down target enhancement, and applied them to singleton-present trials. This cross-condition generalization yielded below-chance decoding at singleton distractor locations. Such below-chance decoding could reflect either enhanced target representations, which would push prediction probabilities for all non-target locations below chance, or suppression specifically targeting the singleton location. To distinguish between these possibilities, we compared prediction probabilities at singleton distractor locations with those at non-singleton distractor locations. Target locations showed above-baseline enhancement beginning around 200 ms, whereas singleton distractor locations showed reliable below-baseline suppression over the same time window. These findings provide converging evidence that singleton distractor representations are actively suppressed within the spatial priority map.

A key limitation of spatial decoding is that target and singleton distractor locations are inherently anticorrelated. Consequently, negative decoding evidence could arise from either suppression of a specific location or enhancement elsewhere. Although our comparison against non-singleton distractor locations partially mitigates this concern, it cannot fully resolve this ambiguity. Future work should test suppression using complementary approaches, such as feature-based decoding or frequency tagging (Forschack et al., 2017; van Moorselaar et al., 2020), particularly in paradigms where target and distractor features vary independently.

We hypothesized that suppression is implemented by coding singleton distractors in an inverted representational format relative to targets. Such inverted coding would facilitate downstream integration of target enhancement and distractor suppression within a shared neural substrate. Consistent with this hypothesis, correlations between target and singleton distractor representations revealed robust negative relationships beginning around 200 ms, indicating that target and singleton distractor locations were inversely coded within a shared neural space. This shared space likely underlies the computation of spatial priorities, allowing target and distractor signals to exert opposing influences. While these findings support the existence of a common coding framework, further work is needed to more precisely characterize the relationship between target and distractor subspaces and their link to the hypothesized priority map.

A substantial body of behavioral, eye-tracking, electrophysiological, EEG, and fMRI evidence supports successful suppression of salient color singleton distractors (Chang & Egeth, 2019; Cosman et al., 2018; Gaspelin et al., 2015, Gaspelin et al., 2017; Sawaki et al., 2012; Sawaki & Luck, 2010; Stilwell et al., 2022; Vatterott & Vecera, 2012). Electrophysiological recordings in NHPs and fMRI evidence in humans have shown reduced neuronal responses to singleton distractors relative to non-singleton distractors (Cosman et al., 2018; Ipata et al., 2006; Adam & Serences, 2021; Won et al., 2020). These findings have been interpreted as evidence for signal attenuation.

However, recent work challenges a purely attenuation-based account. Neural responses to singleton distractors appear highly heterogeneous in V4, FEF and LIP (Klink et al. 2023; Sapountzis et al. 2025). Moreover, decoding analyses indicate that singleton distractor representations persist rather than vanish, suggesting that they may play an active role in guiding behavior. Together, these findings suggest that singleton distractor suppression may rely on representational transformation rather than uniform attenuation.

The present study extends this work by providing direct evidence for a rapid inversion of singleton distractor representations during visual search. We propose that attentional suppression begins with early sensory registration of salient singleton distractors, reflected in the representations observed between 100 - 200 ms. These early signals do not appear to drive behavior. Instead, suppression involves transforming these signals into an inverted format that downregulates the distractor’s priority within a spatial priority map (Fig. 6). The timing of this transformation coincides with the emergence of target representations, suggesting that it may be driven by top-down feedback supporting target guidance rather than by feedforward attenuation alone. Notably, the later inverted singleton representations predicted faster reaction times, providing functional evidence that this transformation supports efficient behavior.

**Fig. 6.**
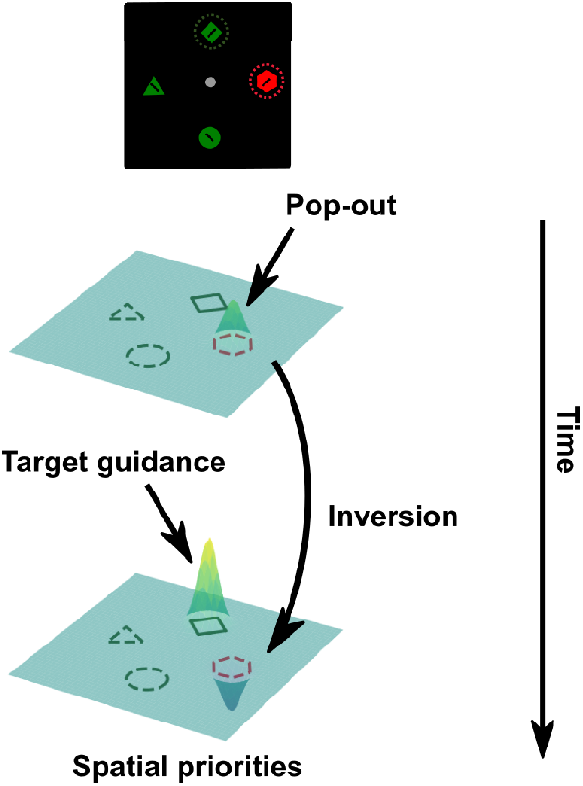
Proposed singleton suppression mechanism within the priority map framework. The process initiates with the registration of singleton distractor information.Subsequently, distractor representations undergo inversion, around the same time as the emergence of target representations. Both distractor and target representations are coded within a shared neural space but in an inverted manner. The representations within this neural space are then accessed to compute the ultimate spatial priorities that guide visual attention.

These findings bear directly on major theoretical accounts of singleton suppression. Both the signal suppression hypothesis (Gaspelin & Luck, 2018b; Sawaki & Luck, 2010) and the rapid disengagement hypothesis (Theeuwes, 2010) posit a transition from initial priority signals to suppression, but differ in the proposed timing. Our results identify a representational transformation beginning around 200 ms. Although the early singleton representations precede this transformation, they were coded in a format distinct from both later singleton representations and target representations. At face value, this pattern could be interpreted as consistent with rapid disengagement hypothesis. However, linking decoding strength to behavior revealed that early singleton representations did not predict reaction times, whereas later representations did. This dissociation suggests that the early distractor representations are unlikely to reflect attentional capture. Instead, these signals may reflect sensory registration of singleton distractors that neither drives attentional capture nor suppression at that early stage of processing.

A limitation of the present study is the absence of eye-tracking data, which precludes directly testing whether early singleton representations reflect covert attentional capture. Nevertheless, several factors argue against this interpretation. First, attentional capture by singleton distractors is rare in similar paradigms, with reported rates typically below 10% (Gaspelin et al., 2017; Stilwell et al., 2023; Adam et al., 2023). Second, when capture does occur, saccade latencies cluster around ∼200 ms (Adam et al., 2023; Gaspelin et al., 2017), coinciding with the onset of the observed transformation. Thus, even if capture occurs, it overlaps temporally with the transformation rather than preceding it. These considerations suggest that the observed representational inversion is unlikely to reflect a purely reactive consequence of attentional capture. Importantly, recent modeling work suggests that the distinction between proactive and reactive suppression may be less fundamental than traditionally assumed. A single mechanism, preventing early distractor signals from triggering an eye movement, can account for both suppression and capture effects depending on relative signal timing (Zhang et al., 2025). Our findings align with this framework by demonstrating how early salience signals can be transformed into suppression-supporting representations.

The two-stage pattern we observed parallels interpretations of ERP components associated with singleton processing. An early positivity (Ppc; 80 - 150 ms) is thought to reflect initial saliency or sensory imbalance, whereas the later Pd (100 - 300 ms) has been linked to inhibitory processing (Barras & Kerzel, 2016; Fortier-Gauthier et al., 2012; Gaspelin et al., 2023; Sawaki & Luck, 2010). Multivariate decoding allows these stages to be distinguished in representational space, providing evidence for a transformation that bridges early saliency signals and later suppression.

Finally, our findings speak to broader theories of spatial priority maps, which integrate goal-driven, experience-driven, and stimulus-driven signals to guide attention (Awh et al., 2012; Fecteau & Munoz, 2006; Luck et al., 2021). Using cross-condition generalization, we found evidence that a common priority representation supports both target enhancement and distractor suppression. However, this interpretation rests on assumptions about shared priority maps that remain debated (Bisley & Mirpour, 2019; Wolfe, 2021). It remains possible that multiple priority maps or representational subspaces coexist, enabling parallel implementation of enhancement and suppression. Disentangling these possibilities remains an important challenge for future work.

In conclusion, we provide evidence for a representational inversion mechanism underlying singleton distractor suppression. Early salience signals are transformed into inverted representations that suppress distractor locations within a shared neural space, supporting efficient visual search.

## Acknowledgements

We would like to acknowledge the invaluable contribution of Dr. Brad Stilwell, Dr. Howard Egeth, and Dr. Nicholas Gaspelin for generously sharing their data. The dataset can be accessed at https://osf.io/9xnsz/.

